# Recapitulation of human embryonic heart beating to promote differentiation of hepatic endoderm to hepatoblasts

**DOI:** 10.1101/2020.05.18.103218

**Authors:** Koki Yoshimoto, Nicholas Minier, Satoshi Imamura, Kaylene Stocking, Janmesh Patel, Shiho Terada, Ken-ichiro Kamei

## Abstract

A microfluidic platform recapitulating human embryonic heart beat improves the functionalization state of hepatocytes derived from hepatic endoderm (HE). Mechanical stretching of mimicked heart beats was applied to HE cells cultured on the microfluidic platform. Stimulated HE-derived hepatoblasts increased cytochrome P450 3A (CYP3A) metabolic activities and hepatoblast functional markers expression, leading for advancement of regenerative medicine and drug screening.

---

Hepatocytes are major components of the liver and have essential physiological roles, such as protein synthesis/storage, glucose metabolism/storage, detoxification, and excretion of exogenous molecules. Disruption of hepatic functions causes severe problems, such as hepatic cirrhosis and liver cancers, resulting patient death.^1,2^ Liver transplantation is the only method for curing these patients. Moreover, drug discovery requires the use of primary hepatocytes to evaluate the toxicity of drug candidates prior to clinical trials.^3^ In both cases, it remains problematic to identify the suitable healthy donors to obtain livers or hepatocytes. There is an urgent need to develop alternatives to hepatocytes.

To fulfil these requirements, human pluripotent stem cells (hPSCs), such as human embryonic and induced pluripotent stem cells (hESCs^4^ and hiPSCs^5,6^, respectively), show high potential because of their capability for unlimited self-renewal and differentiation to almost any type of tissue cells. However, although many studies have been performed to obtain hPSC-derived hepatocytes, they remain immature as hepatocytes, showing fewer functional properties. Existing differentiation protocols mostly used biochemical factors, such as fibroblast growth factor (FGF) and bone morphogenesis protein (BMP). Although such biochemical factors have been investigated intensively, the effects of biomechanical forces on the hepatic developmental process remain unknown. As mechanical forces regulate a variety of biological contexts, including molecules, cells, tissues, and organs^7,8^, their effects must be considered to update hepatic differentiation methods from hPSCs.

In this regard, the hepatic endoderm (HE) is the critical stage affected by mechanical forces. The HE is formed at early developmental stages from the definitive endoderm, and HE-derived hepatoblasts give rise to hepatocytes or cholangiocytes. Notably, the HE is exposed to oscillating mechanical forces because of the heart beating (Fig. 1a). Although biochemical factors have been reported in static culture conditions^9,10^, the effects of mechanical forces that induce differentiation of the HE into hepatoblasts has not been investigated because of limited access to human embryos and the lack of proper *in vitro* models that recapitulate the *in vivo* physiological embryonic developmental process. Therefore, current protocols cannot be used when evaluating mechanical forces from embryonic heart beats, resulting immature differentiation towards hepatocytes.

**Fig. 1.**
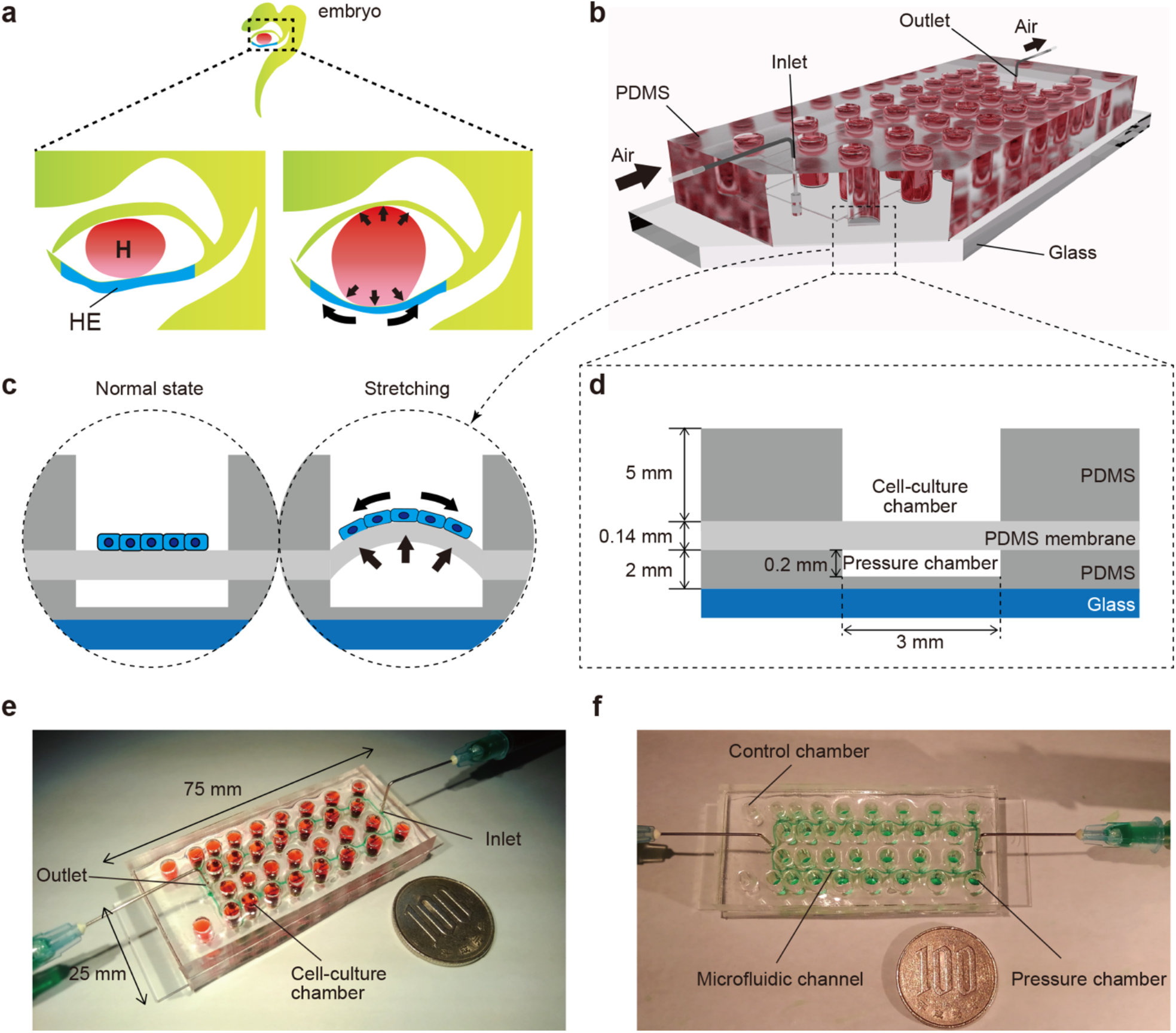
**a**, Illustration of early human embryo. Heart (H: red) beating confers mechanical stimulation to the surrounding cells, and hepatic endoderm (HE: blue) which differentiates to hepatoblasts, is exposed to mechanical forces. **b,c**, Appearance (**b**) and cross-section (**c**) of a microfluidic device for applying a series of stretching stimulations to HE cells (HECs). **d**, Cross-section view of the device, which is composed of polydimethylsiloxane (PDMS) and consists of a top layer with cell-culture chambers, middle membrane layer, and bottom layer with pressure chambers on a glass slide. **e**, Photograph of actual our device fabricated on a glass slide (25 × 75 mm). Culture chambers is filled with red ink. This device has two sets of culture chambers in which cells are cultured under the same intensity of mechanical stimulation. **f**, Photograph of a microfluidic device. Microfluidic channels and pressure chambers filled with green ink.

Microfluidic technology shows potential for applying mechanical forces to cells, as it allows for systematic manipulation of the cell culture conditions (e.g., flow dynamics, cell-cell/matrix interactions, and mechanical stretching) in two- and three-dimensional manners, which cannot be achieved using conventional cell-culture models. Recently, organs-on-a-chip platforms based on microfluidic technology have been reported to recapitulate physiological mechanical forces *in vitro* using natural tissues^11–14^. However, most organ-on-a-chip platforms can stimulate cells using only a single mechanical condition^15,16^, and thus the optimal mechanical strength for obtaining targeted functional cells cannot be determined.

Here, we developed a microfluidic platform for applying multiple mechanical forces on hPSC-derived HE cells to identify the optimal mechanical stress that facilitates the differentiation of HE cells to functional hepatoblasts (**Fig. 1b**). We developed a microfluidic device composed of polydimethylsiloxane (PDMS) elastic material with a ballooned thin membrane as the cell-culture substrate which can be actuated to mimic heart beats in an embryo. The balloons with cells inflate and deflate repeatedly via pressure regulation (**Fig. 1c and d**). We showed that hPSC-derived hepatoblasts differentiated under the optimal stretching condition with the expression of drug metabolism enzymes and proteins specific to hepatoblasts. These findings demonstrate that the dynamic mechanical forces are critical for differentiation from the HE to hepatoblasts.

To recreate the embryonic heart beat *in vitro*, a microfluidic device with a series of stretchable balloon membranes was fabricated (**Fig. 1b,e,f and ESI Fig. S1**). This microfluidic device consisted of three layers: a top layer for cell-culture wells, middle layer of thin membrane as the stretchable cell-culture substrate, and bottom layer for forming pressure chambers. The top layer is 5 mm thick, and each well in the top layer is 3 mm in a diameter. The middle PDMS membrane is 0.14 mm thick. The bottom layer is 2 mm thick with a 0.25-mm channel and chamber height and 0.2-mm channel width (**ESI Fig. S2**). The molds for the top and bottom layers were fabricated with a high-resolution 3D printer^17^.

The thin ballooned PDMS membrane was actuated with the regulator connected to an air compressor. To test a series of stretching forces within a single device, we used a pressure-drop method^18^ in which air pressures was decreased in an inverse proportion to the length of fluidic flow (**Fig. 2a**). The ¥ pressure drop (Δ*P*) for incompressible fluid flow was determined from the Fanning friction factor (f) using the Fanning formula:

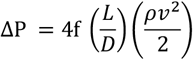

Hydraulic diameter, D (m) was calculated as:

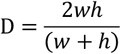

where *w* and *h* represent the channel width and height, respectively, *L* is the channel length, ρ is the fluid density, and *v* is the average velocity in a channel. The amount of air pressure applied to each chamber decreased with increasing channel length. The device was designed to have 2 sets of 15 culture chambers along a micro channel and negative control chambers in a single device.

**Fig. 2.**
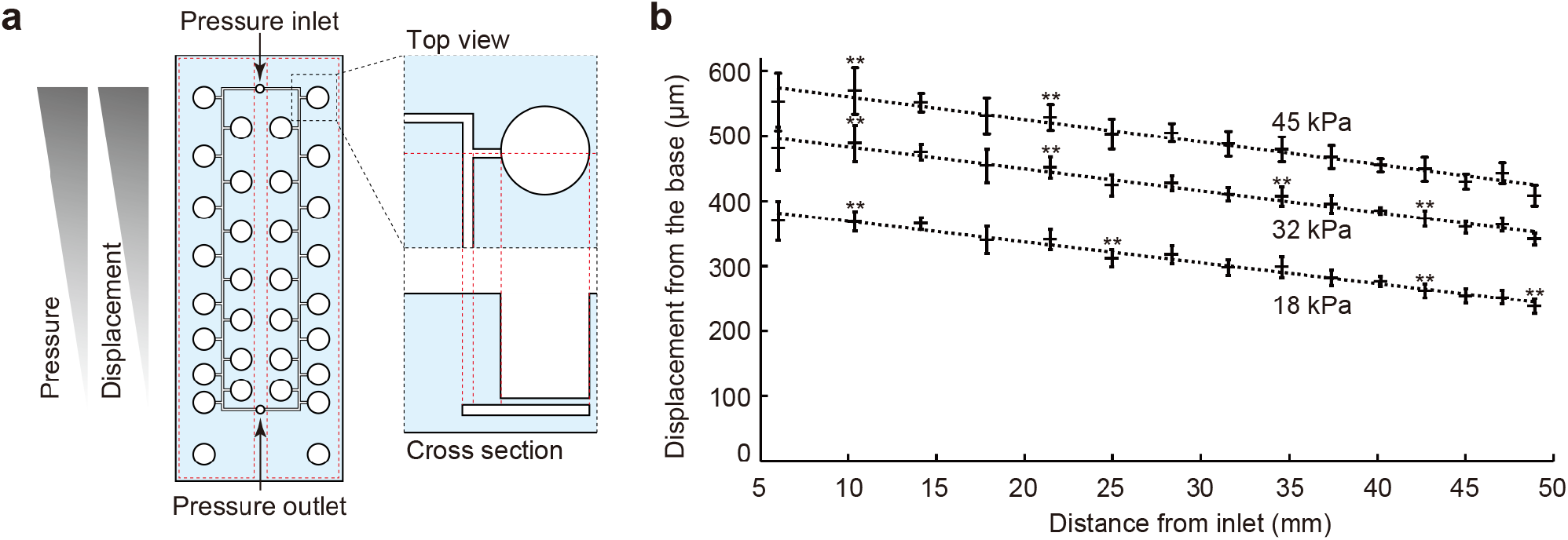
Pressure-drop method to generate a series of PDMS membrane displacements in a single device. **a**, Membrane displacements are inversely proportional to the distance from the inlet because of a pressure drop along with a microfluidic channel. **b**, Displacement measurement of the device with CCD laser displacement camera when 18, 32, and 45 kPa were applied at the inlet. ANOVA with Tukey-Kramer test compared with all displacements of the pressure chambers at 18, 32, and 45 kPa: \*\**P* < 0.01. Each plot represents the mean ± standard deviation determined from three independent experiments measuring the two chambers in a single device.

To demonstrate the pressure-drop method over a series of membrane stretching events, we applied three input pressures (i.e., 18, 32, and 45 kPa) to the inlet and measured the vertical displacement from the base of the membrane (**Fig. 2a**). As expected, membrane displacements corresponded to pressure decreases for the tested input pressures. When more than 45 kPa pressure was applied to the inlet, our device was unstable because of air leakage. Based on these results, we selected 11 different displacements and labeled the input pressure (kPa) as the distance from the inlet (mm)], such as (45, 10.37) (**Fig. 2b**).

Then, hPSCs were differentiated to hepatoblasts in a device (**Fig. 3a** and see in ESI)^19,20^. Prior to cell culture in a device, the cell-culture chambers were coated with Matrigel. The differentiation from hPSCs to hepatoblasts into three stages. Briefly, in the first stage, hPSCs were directed into definitive endoderm (DE) by treatment with 100 ng mL^−1^ activin A, 10 μM ROCK inhibitor, 3 μM CHIR99021, 10 μM LY294002, 10 ng mL^−1^ BMP4, and 100 ng mL^−1^ basic FGF (bFGF). To confirm DE differentiation, expression of CXCR4 (CXC chemokine receptors), a DE cell-surface marker, was evaluated by flow cytometry (**Fig. 3b**). More than 99% of cells expressed CXCR4, indicating efficient DE differentiation from hPSCs. In the second stage, DE cells were treated with a lower concentration of activin A (50 ng μL^−1^) to obtain HECs. In the third stage, HECs were differentiated to hepatoblasts by treatment with 20 ng mL^−1^ BMP4 and 10 ng mL^−1^ FGF10. Stretching stimulation at 0.2 Hz was applied during the third stage for 4 days. Cells were observed on the PDMS thin membrane by day 12 under conditions without mechanical forces (**ESI Fig. S3**).

**Fig. 3.**
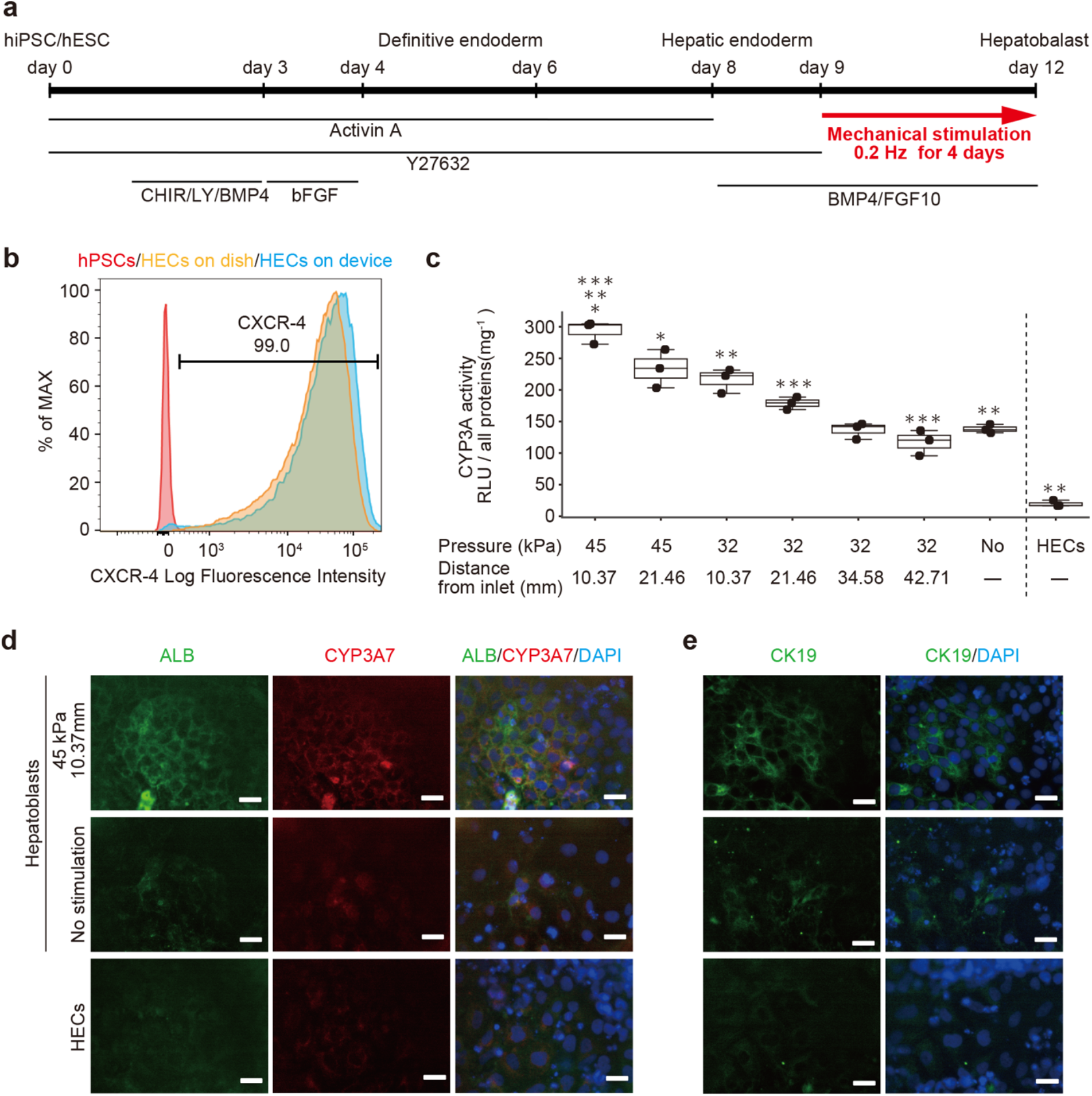
Hepatoblast differentiation from human pluripotent stem cells (hPSCs) promoted by mechanical forces on hepatic endoderm-like cells (HECs). **a**, Schematic diagram showing hepatic differentiation from hPSCs. ROCK inhibitor (Y27632), WNT inhibitor (CHIR, CHIR99021), PI3K inhibitor (LY, LY294002), bFGF, BMP4, and FGF10 were used for corresponding differentiation stages (see in ESI). **b**, Flow cytometric analyses showing the proportion of CXCR4 expression in hPSCs, HECs in a dish and HECs in a microfluidic device. **c**, Bioluminescent CYP3A activity assay for HECs and HBCs. ANOVA with Tukey-Kramer test compared with relative light units (RLU) of all samples: *P < 0.05. **P and ***P < 0.01. Max, median, minimum of three independent experiments were shown. **d and e**, Immunocytochemical analyses showing the expression of ALB, CYP3A7, (**d**) and CK19 (**e**) in HECs and HBCs in the indicated conditions. Nuclei were stained with DAPI. Scale bars represent 50 μm.

To investigate the effect of mechanical forces on the differentiation of HECs to HBCs, the activities of cytochrome P450 3A (CYP3A), which was specifically expressed in hepatic cells, was measured in a bioluminescent CYP3A activity assay at day 12 (**Fig. 3c and see ESI**). Compared with HECs, non-stimulated hepatoblasts showed significantly higher activities, as expected. When 32-kPa input pressure was applied to the larger displacements at (32, 21.46) and (32, 10.37), stimulated hepatoblasts showed higher CYP3A activities, whereas CYP3A activities at (32, 42,71) and (32, 34.58) did not show dramatic differences compared with non-stimulated HECs. Moreover, at a higher input pressure of 45 kPa, the larger displacements at (45, 21.46) and (45, 10.37) gave significantly higher CYP3A activities in hepatoblasts, and hepatoblasts at (45, 10.37), which were at least two-fold higher CYP3A activities. These results suggest that mechanical stimulation by stretching cell-culture substrates increase the hepatoblastic metabolic activities.

To further investigate the effects of stretching stimulation during HEC-hepatoblast differentiation, the protein expression of albumin (ALB), CYP3A7, and cytokeratin 19 (CK19), which are specifically expressed in hepatoblasts^21,22^, were observed by immunocytochemistry (Fig. 3d for ALB and CYP3A7, and Fig. 3e for CK19). Hepatoblasts at (45, 10.37) showed higher expression of ALB, CYP3A7, and CK19 proteins than non-stimulated hepatoblasts, which agreed with the results observed for CYP3A activities. HECs did not express the tested proteins. Generally, the tested proteins are expressed *in vivo* hepatoblasts but not *in vitro*. These results suggest that the stretching stimulation makes hPSC-derived hepatoblasts more functional than those in conventional cell-culture.

## Conclusions

In summary, we developed a microfluidic device by applying multiple stretching stimulation during HEC-hepatoblast differentiation from hPSCs. Using this device, we found that mechanical stimulation improved the functionalities of hepatoblasts. Although the underlying mechanisms should be investigated to further improve mechanically stimulated HEC-hepatoblast differentiation, previous reports of the effects of mechanical stimulation to epithelial,^23^ endothelial,^8,24^ and mesenchymal stem cells^25–27^ suggest that mechanical stimuli via cell-substrate interactions influence the translocation of the transcription factor yes-associated protein/transcriptional coactivator with the PDZ-binding motif in the Hippo pathway. Our device and approach can provide not only insight into the hepatic developmental process but also tools for applications in both drug discovery and regenerative medicine.

## Supporting information

Supplemental informations

## Acknowledgements

Funding was generously provided by the Japan Society for the Promotion of Science (JSPS; 16K14660, 17H02083, 18KK0306, and 19H02572), Japan Agency for Medical Research and Development (AMED; 17937667) and LiaoNing Revitalization Talents Program (XLYC1902061). NM was supported by BME Paris Master program. KS and JP were supported by the Nakatani Foundation for advancement of measuring technologies in biomedical engineering. The WPI-iCeMS is supported by the World Premier International Research Centre Initiative (WPI), MEXT, Japan.

## Author Contributions

Conceptualization: KY NM KK, Data curation: KY NM SI KS JP ST, Formal Analysis: KY NM, Funding acquisition: KK, Investigation: KY NM KK, Methodology: KY NM KK, Project administration: KK, Resources: KK, Software: KK, Supervision: KK, Validation: KY KK, Visualization: KY KK, Writing – original draft: KY KK, Writing – review & editing: KY NM KS JP KK.

