## Supplemental informations for "Recapitulation of human embryonic heart beating to promote differentiation of hepatic endoderm to hepatoblasts"

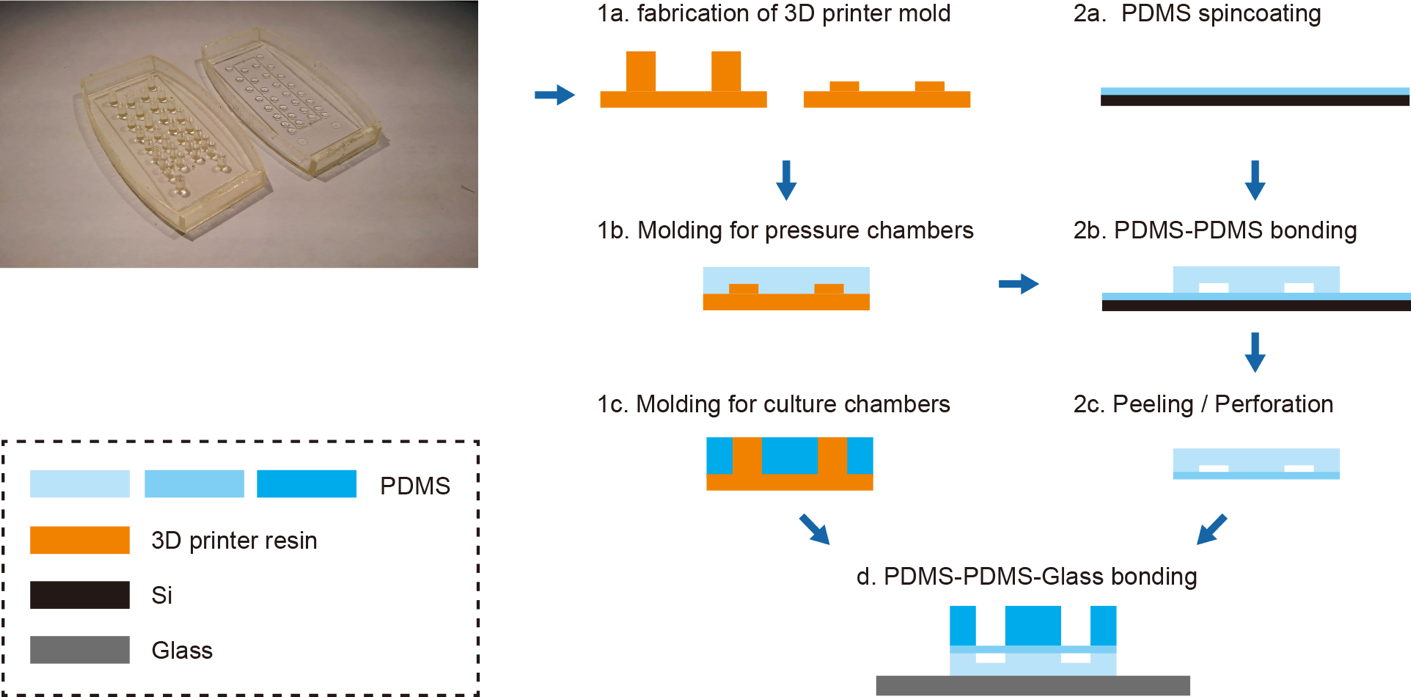

**Supplementary Fig. S1.** Fabrication of a microfluidic device that recapitulates human embryonic heart beating. The device was fabricated as follows: (1a) For the top and bottom layer, the molds were fabricated with a 3D printer. (1b,c) PDMS casted into the molds and cured at 80°C for over 16 h. (2a) PDMS was dropped on the silicon wafer, spin-coated at 500 rpm for 30 s, and cured at 80°C for 10 min. (2b) The top layer and PDMS membrane were bonded together and baked at 80°C for over 1.5 h. (2c) The PDMS-PDMS forms were peeled off (d) and bonded to the top layer and glass.

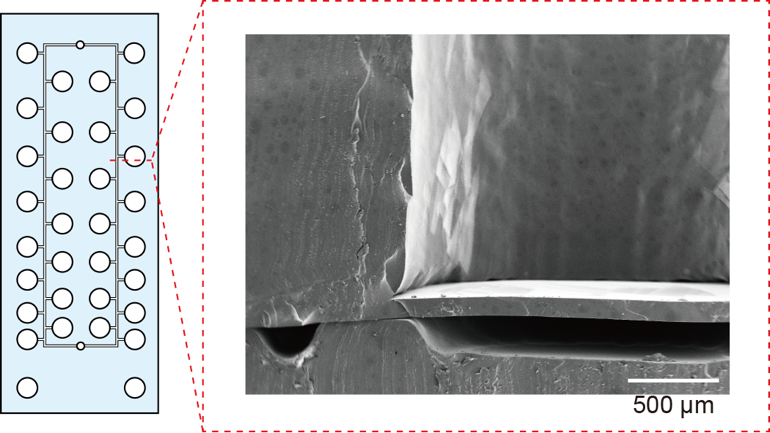

**Supplementary Fig. S2. Scanning electron** micrograph of cross-section in the microfluidic device. The micro channel and pressure chamber are indicated. Scale bar, 500 μm.

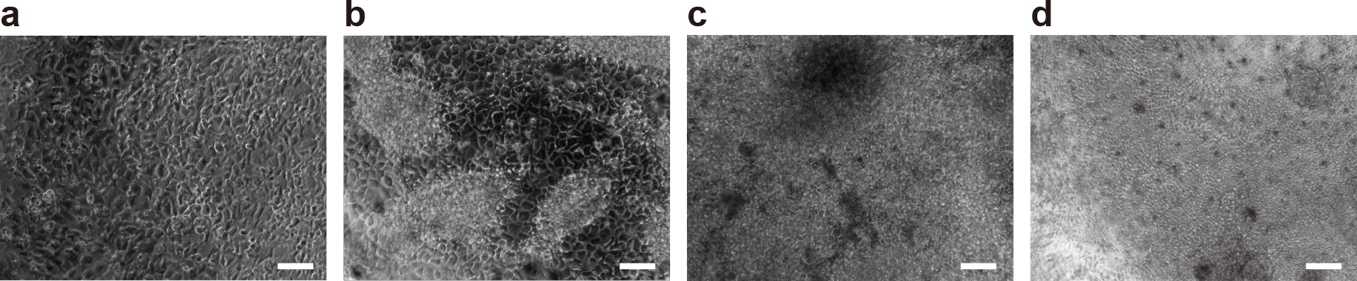

**Supplementary Fig. S3.** Microphotograph of hPSCs-derived cells. **a,** hPSC-derived definitive endoderm (DE) cells at day 1. **b,** DE cells at day 4. **c,** Hepatic endoderm (HE) cells at day 8. **d,** HE-derived hepatoblasts. Scale bars represent 100 μm.

**Materials and Methods**

**Chip design.** The formula of a pressure drop and length in microfluidic channel reported by Shimizu *et al.* was adapted.^1^ The mold of pressure chambers and culture chambers were designed with a specific channel height (200 μm) and width (100 μm), and specific branch channel length (1 mm) by computer-aided design. The culture chambers were replaced considering the pressure drops.

**Microfluidic device fabrication.** A microfluidic device was fabricated using stereolithographic 3D-printing techniques and solution cast-molding processes.^2^ The molds for the top layer and bottom layer were produced using a 3D printer (Keyence Corporation, Osaka, Japan). After fabricating the molds with the 3D printer, the molds were washed with 99.9% EtOH for over 12 h. The molds were dried at 80°C for over 30 min. Sylgard 184 PDMS two-part elastomer (10:1 ratio of pre-polymer to curing agent; Dow Corning Corporation, Midland, MI, USA) was mixed, poured into a 3D-printed mold to produce a 5-mm-thick PDMS top layer and a 2-mm-thick PDMS bottom layer, and de-gassed by using a vacuum desiccator for over 30 min. The PDMS material was then cured in an oven at 80°C for over 16 h. After curing, the PDMS form was removed from the mold, trimmed, and cleaned. Sylgard 184 PDMS two-part elastomer (10:1 ratio of pre-polymer to curing agent) was mixed, dropped onto the silicon wafer, spin-coated at 500 rpm for 30 s, and baked at 80°C for 10 min. The pressure chamber layer and PDMS thin membrane on silicon wafer were corona-plasma-treated (Kasuga Denki, Inc., Kawasaki, Japan), bonded together, baked at 80°C for 1.5 h, and peeled off from the silicon wafer. The top layer-thin membrane forms, bottom layer, and glass were corona-plasma-treated and bonded together by baking in an oven at 80°C for 18 h. The devices were used within one day of completion.

**Scanning electron micrograph (SEM).** A 5-nm-thick platinum film was deposited on the device sectioned with a knife by sputtering (MSP 30T; Shinku Device, Ibaraki, Japan). Images of the device were acquired with a scanning electron microscope at 10 kV (JCM-5000; JEOL Ltd., Tokyo, Japan).

**Device control.** Air flow was controlled by an air regulator. The regulator was controlled by the LabVIEW program (National Instruments, Austin, TX, USA).

**Measurement of pressure to inlet and PDMS displacement.** The pressure was measured close to the inlet with a pressure sensor (ZS-46-5F; SMC, Tokyo, Japan). Vertical displacement of the PDMS thin membrane was measured with a CCD laser displacement sensor (LK-G5000; Keyence).

**hPSC culture.** hESCs were used according to the guidelines of the ethics committee of Kyoto University. H9 hESCs were purchased from WiCell Research Institute (Madison, WI, USA). H9 hESCs were used in this study. Prior to culture, hESC-certified Matrigel (Corning, Inc., Corning, NY, USA) was diluted with Dulbecco’s modified Eagle medium (DMEM)/F12 (Merck KGaA, Damstadt, Germany ) at a 1:75 (v/v) ratio and coated onto a culture dish. Matrigel was incubated in a dish for 24 h at 4°C. Excess Matrigel was removed, and the coated dish was washed with fresh DMEM/F12.

mTeSR-1-defined medium (Stem Cell Technologies, Vancouver, Canada) supplemented with 1% (v/v) penicillin/streptomycin (Fujifilm Wako, Osaka, Japan) was used for daily culturing of hPSCs. For passaging, the cells were dissociated with TrypLE Express (Thermo Fisher Scientific, Waltham, MA, USA) for 3 min at 37°C and harvested. A cell strainer was used to remove undesired cell aggregates from the cell suspension, and the cells were centrifuged at 200 ×*g* for 3 min and resuspended in mTeSR-1 medium. The cells were counted using Via 1-Cassete^TM^ (ChemoMetec A/S, Gydevang, 43, Denmark) of a NucleoCounter NC-200 (Chemetec, Baton Rouge, LA, USA). mTeSR-1 medium containing 10 µM of the ROCK inhibitor Y-27632 (Fujifilm Wako) was used to prevent apoptosis of dissociated hPSCs on day 1. mTeSR-1 medium without ROCK inhibitor was used on subsequent days, with daily medium changes.

**Hepatic differentiation from hPSCs on device.** Prior to inducing differentiation, the culture chambers of the device were coated with Matrigel at 35°C for 60 min. Matrigel was removed with an aspirator.

To induce endoderm differentiation, cultured hPSCs were washed with D-PBS (no calcium, no magnesium) (Thermo Fisher Scientific) and treated with TryPLE Express at 37°C for 3 min, followed by addition of basal medium and transfer of the cell suspension into a 15-mL tube. Cells were centrifuged at 200 ×*g* for 3 min and the supernatant was removed. The cells were resuspended to 7.00 × 10^5^ cells mL^-1^ in mTeSR-1 medium supplemented with 10 µM Y27632 and 100 ng mL^−1^ activin A (human recombinant) (Fujifilm Wako), applied 30 μL chambers^-1^ resuspended solution to a Matrigel-coated culture chambers, and cultured in a humidified incubator at 37°C with 5% CO_2_ for 24 h. At the end of day 1, the medium was replaced with fresh mTeSR-1 medium supplemented with 10 µM Y27632 and 100 ng mL^−1^ activin A and cultured for another 24 h. On day 2, the medium was replaced with mTeSR-1 medium supplemented with 10 µM Y27632, 100 ng mL^−1^ activin A, 10 ng mL^−1^ BMP-4 (human recombinant) (R&D Systems, Minneapolis, MN, USA), 10 µM LY294002 (Cayman Chemical, Arbor, MI, USA), and 3 µM CHIR99021(ReproCELL, Kanagawa, Japan), and the cells were incubated for 24 h. On day 3, the medium was replaced with mTeSR-1 medium supplemented with 10 µM Y27632, 100 ng mL^−1^ activin A, 10 ng mL^−1^ BMP-4, and 10 µM LY294002, and the cells were incubated for 24 h. On day 4, the medium was replaced with Roswell Park Memorial Institute 1640 (RPMI) medium, GlutaMax^TM^ Supplement (Thermo Fisher Scientific), supplemented with 2% (v/v) B-27^TM^ supplement (Thermo Fisher Scientific), 1% (v/v) MEM Non-essential Amino Acid Solution without L-glutamine, liquid, sterile-filtered Bioreagent, suitable for cell culture (NEAA) (Merck KGaA), 1% (v/v) penicillin/streptomycin, 10 µM Y27632, 100 ng mL^−1^ activin A, and 100 ng mL^−1^ bFGF (human recombinant) (Fujifilm Wako), and the cells were incubated for 24 h.

To induce hepatic endoderm specification, the cells were treated with RPMI medium GlutaMax^TM^ Supplement, supplemented with 2% (v/v) B-27^TM^ supplement, 1% (v/v) NEAA, 1% (v/v) penicillin/streptomycin, 10 µM Y27632 and 50 ng mL^−1^ activin A, with daily medium changes for 3 days. On day 8, to induce hepatoblast specification, the cells were treated with RPMI medium GlutaMax^TM^ Supplement, supplemented with 2% (v/v) B-27^TM^ supplement, 1% (v/v) NEAA, 1% (v/v) penicillin/streptomycin, 25 mM HEPS (Fujifilm Wako), 10 µM Y27632, 20 ng mL^−1^ BMP-4 and 10 ng mL^−1^ FGF-10 (human recombinant) (R&D Systems). The cells were then treated with RPMI medium GlutaMax^TM^ Supplement, supplemented with 2% (v/v) B-27^TM^ supplement, 1% (v/v) NEAA, 1% (v/v) penicillin/streptomycin , 25 mM HEPES, 20 ng mL^−1^ BMP-4 and 10 ng mL^−1^ FGF-10, with daily medium changes and 0.2 Hz mechanical stimulation for 4 days.

**Definitive endoderm differentiation on dish for flow cytometry.** Prior to inducing differentiation, a cell-culture dish was coated with 0.1% gelatin from porcine skin, type A (Merck KGaA) in phosphate-buffered saline (PBS) at 25°C room temperature for 30 min. The gelatin solution was then aspirated and DMEM/F12 medium supplemented with 10% (v/v) fetal bovine serum (JRH Biosciences, St, Lenexa, KS, USA), 1% (v/v) L-glutamine, 200 mM Solution (Thermo Fisher Scientific), 1% (v/v) penicillin/streptomycin, and 100 µM β-mercaptoethanol (Fujifilm-Wako) was introduced onto the culture dish for serum coating at 37°C for 24 h. The coated dish was then rinsed with fresh medium.

To induce endoderm differentiation, cultured hPSCs were washed with PBS and treated with TryPLE Express at 37°C for 3 min, followed by addition of basal medium and transfer of the cell suspension into a 15-mL tube. The cells were centrifuged at 200 ×*g* for 3 min, after which the supernatant was removed. The cells were resuspended in mTeSR-1 medium supplemented with 1% (v/v) penicillin/streptomycin, 10 µM Y27632 and 100 ng mL^−1^ activin A, plated on a serum-coated culture dish, and cultured in a humidified incubator at 37°C with 5% CO_2_ for 24 h. At the end of day 1, the medium was replaced with fresh mTeSR-1 medium supplemented with 1% (v/v) penicillin/streptomycin, 10 µM Y27632 and 100 ng mL^−1^ activin A and cultured for another 24 h. On day 2, the medium was replaced with mTeSR-1 medium supplemented with 1% (v/v) penicillin/streptomycin, 10 µM Y27632, 100 ng mL^−1^ activin A, 10 ng mL^−1^ BMP-4, 10 µM LY294002, and 3 µM CHIR99021, and cells were incubated for 24 h. On day 3, the medium was replaced with mTeSR-1 medium supplemented with 1% (v/v) penicillin/streptomycin, 10 µM Y27632, 100 ng mL^−1^ activin A, 10 ng mL^−1^ BMP-4, and 10 µM LY294002, and the cells were incubated for 24 h. On day 4, medium was replaced with RPMI medium, GlutaMax^TM^ Supplement, supplemented with 2% (v/v) B-27^TM^ supplement, 1% (v/v) NEAA, 1% (v/v) penicillin/streptomycin, 100 ng mL^−1^ activin A, and 100 ng mL^−1^ bFGF, and the cells were incubated for 24 h. The cells were treated with RPMI medium, GlutaMax^TM^ Supplement, supplemented with 2% (v/v) B-27^TM^ supplement, 1% (v/v) NEAA, 1% (v/v) penicillin/streptomycin, and 50 ng mL^−1^ activin A, with daily medium changes for 3 days. The cells were then treated with RPMI medium, GlutaMax^TM^ Supplement, supplemented with 2% (v/v) B-27^TM^ supplement, 1% (v/v) NEAA, 1% (v/v) penicillin/streptomycin, 20 ng mL^−1^ BMP-4, and 10 ng mL^−1^ FGF-10, with daily medium changes for 2 days. On day 6, the cells were harvested for flow cytometry.

**Flow cytometry.** The cells were harvested with TrypLE Express and rinsed with PBS twice prior to cell counting. For staining with antibodies, the cells were diluted to a final concentration of 1 × 10^7^ cells mL^−1^ in stain buffer (fetal bovine serum) (BD Pharmingen, Franklin Lakes, NJ, USA). Fluorescence-labeled antibodies (APC Mouse Anti-Human CD184 (CXCR4), clone 12G5; BD Pharmingen) were added and incubated at room temperature for 1 h. As a negative control, specific isotype controls (APC Mouse IgG2a k Isotype Control, clone G155-178; BD Pharmingen) were used. After removing excess antibodies by centrifugation at 300 ×*g* for 5 min, the cells were washed with staining buffer, and cell suspensions were applied to a FACS Canto II (BD Biosciences, Franklin Lakes, NJ, USA) for flow cytometric analysis. Data analysis was performed using FlowJo software (v9; FlowJo, LLC, Ashland, OR, USA).

**Cytochrome P450-GloTM assays with luciferin.** To perform cytochrome P450-GloTM assays with luciferin (Promega, Madison, WI, USA), 2/3 medium, 20 mL medium was removed. Proluciferin IPA was diluted to RPMI medium, GlutaMax^TM^ Supplement, supplemented with 2% (v/v) B-27^TM^ supplement, 1% (v/v) NEAA, and 1% (v/v) penicillin/streptomycin. (1.5:1000). The air flow, mechanical stimulation was stopped, and 20 mL medium containing proluciferin IPA was applied to the culture chambers. The cells were incubated at 37°C with 5% CO_2_ for 1 h. Next, 25 mL ×2 medium containing luciferin IPA was collected from two culture chambers with the same mechanical stimulation and added to 96-well plates (White Microwell SI; Thermo Fisher Scientific); 50 μL P450-Glo^TM^ Reagent was added, incubated at 28°C for 20 min, and relative light units were measured with a Synergy HTX Microplate Reade (Biotek, Winooski, VT, USA) at 28°C.

**Collection of all proteins in cells.** The cells were rinsed with D-PBS (-) (Fujifilm Wako) 3 times, harvested with Tryple Express, and collected into 1.5-mL tubes. Next, 0.5 mL RPMI medium, GlutaMax^TM^ Supplement, supplemented with 2% (v/v) B-27^TM^ supplement, 1% (v/v) NEAA, and 1% (v/v) penicillin/streptomycin was added. The tube was centrifuged at 3000 ×*g* for 3 min, and the supernatant was removed. The cells were rinsed with cold D-PBS and 100 μL cold 1× RIPA buffer (Cell Signaling Technology, Danvers, MA, USA) in double distilled water was added, mixed with a vortex mixer, and incubated on ice for 30 min. The cells were sonicated at 4°C with a sonicator (US-1R cleaner; AS ONE, Osaka, Japan), and centrifuged at 4°C, 10,000 ×*g* for 20 min. The supernatant was collected into new 1.5-mL tubes and stored at -20°C.

**Quantification of all proteins in cells.** Using a BCA protein kit (TAKARA, Shiga, Japan), working solution (BCA reagent A: B = 100:1) and 0.2 mg μL^-1^ BCA standard were mixed with the collected proteins in 100 μL 1× RIPA buffer in a 96-well plate (Matrix Microplate w/lids 96-well blk/clr, flat bottom, tissue culture, PS; Thermo Fisher Scientific). Proteins were incubated at 37°C for 60 min and measured at a wavelength of 562 nm with a Synergy HTX Microplate Reader.

**Immunocytochemistry.** The cells were fixed with 4% paraformaldehyde in D-PBS (-) (Fujifilm Wako) for 20 min at 25°C and then permeabilized with 0.5% (v/v) Triton X-100 in D-PBS for 16 h at 25°C. Subsequently, the cells were blocked in D-PBS (5% (v/v) normal goat serum blocking solution (Maravai Life Sciences, San Diego, CA, USA), 5% (v/v) normal donkey serum (Jackson ImmunoResearch, West Grove, PA, USA), 3% (v/v) albumin, essentially globulin-free (Merck KGaA), and 0.1% Tween-20 (Nacalai Tesque, Kyoto, Japan) at 4°C for 16 h and then incubated at 4°C for 16 h with the primary antibody [anti-human albumin mouse IgG, 1:500; R&D Systems: anti-human cytokeratin 19 (CK19) mouse IgG, 1:500; Thermo Fisher Scientific: anti-human cytochrome P450 3A7 (CYP3A7) rabbit IgG, 1:500; Proteintech, Chicago, IL, USA] in DPBS with 0.5% Triton X-100 (MP Biomedicals, Santa Ana, CA, USA). The cells were then incubated at 37°C for 60 min with a secondary antibody (AlexaFluor 488 Donkey anti-mouse IgG (H+L), 1:1000; Jackson ImmunoResearch: AlexaFluor 594 Donkey anti-rabbit IgG (H+L) , 1:1000) in blocking buffer prior to a final incubation with 4′,6-diamidino-2-phenylindole (D212 -Cellstain®- DAPI) (Fujifilm Wako) at 25°C for 30 min.

**Image acquisition**. The sample containing cells was placed on the stage of a Nikon ECLIPSE Ti inverted fluorescence microscope equipped with a CFI plan fluor 10×/0.30 N.A. objective lens (Nikon, Tokyo, Japan), CCD camera (ORCA-R2; Hamamatsu Photonics, Hamamatsu City, Japan), mercury lamp (Intensilight; Nikon), XYZ automated stage (Ti-S-ER motorized stage with encoders; Nikon), and filter cubes for fluorescence channels (DAPI, GFP HYQ, TRITC; Nikon). For image acquisition, the exposure times were set to 200 ms for DAPI, 200 ms for GFP HYQ (for ALB), 50 ms for GFP HYQ (for CK19), and 200 ms for TRITC (for CYP3A7).
